# Elevational and Oceanic Barriers Shape the Distribution, Dispersal and Diversity of Aotearoa’s Kapokapowai (*Uropetala*) Dragonflies

**DOI:** 10.1101/2024.11.25.625207

**Authors:** Ethan R. Tolman, Christopher D. Beatty, Aaron Goodman, Priscilla Ramchand, Tara McAllister, Kelly Reyes, Akifa Zahara, Kassandra Taveras, Vincent Wade, John Abbott, Seth Bybee, Robert Guralnick, Kathleen M. Harding, Manpreet K. Kohli, Paul B. Frandsen, Jessica Ware, Anton Suvorov

## Abstract

Mountains and islands provide an opportunity for studying the biogeography of diversification and population fragmentation. Aotearoa (New Zealand) is an excellent location to investigate both phenomena due to alpine emergence and oceanic separation. While it would be expected that separation across oceanic and elevation gradients are major barriers to gene flow in animals, including aquatic insects, such hypotheses have not been thoroughly tested in these taxa. By integrating population genomic from sub-genomic Anchored-Hybrid Enrichment sequencing, ecological niche modeling, and morphological analyses from scanning-electron microscopy, we show that tectonic uplift and oceanic vicariance are implicated in speciation and population structure in Kapokapowai (*Uropetala*) dragonflies. Although Te Moana o Raukawa (Cook Strait), is likely responsible for some of the genetic structure observed, speciation has not yet occurred in populations separated by the strait. We find that the altitudinal gradient across Kā Tiritiri-o-te-Moana (the Southern Alps) is not impervious but it significantly restricts gene flow between aforementioned species. Our data support the hypothesis of an active colonization of Kā Tiritiri-o-te-Moana by the ancestral population of Kapokapowai, followed by a recolonization of the lowlands. These findings provide key foundations for the study of lineages endemic to Aotearoa.

## Introduction

Islands are widely recognized for their unique role in shaping biodiversity (Barreto et al., 2021; Conway & Olsen, 2019; Gleditsch et al., 2022; Wilson, 2009). The Theory of Island Biogeography by MacArthur and Wilson (1963) summarized several of these phenomena, and this theory was later extended to mountaintop habitats (Wyckhuys et al., 2022). The islands of Aotearoa (New Zealand; see table 1 for nomenclature) have often been considered as a natural laboratory to study both island and mountain biogeography (Shepherd et al., 2024; Thomas et al., 2023; Wallis & Buckley, 2024; Waters & Craw, 2006). This is due to the opening and closing of Te Moana o Raukawa (Cook Strait) which separates Te Ika-a-Māui and Te Waipounamu (also known as the North and South Islands respectively) and last opened 500,000 years ago (Lewis et al., 1994), and the uplift of Kā Tiritiri-o-te-Moana (the Southern Alps), a relatively young mountain range on Te Waipounamu (South Island; Table 1).

**Table 1:** Reference table of colonial and Mãori names of localities and taxa.

| **Geographic localities** |  |
| --- | --- |
| <u>Māori name</u> | <u>Colonial name</u> |
| Aotearoa | New Zealand |
| Te Moana o Raukawa | Cook Strait |
| Te Ika-a-Māui | North Island |
| Te Waipounamu | South Island |
| Kā Tiritiri-o-te-Moana | Southern Alps |
| **Population Names** |  |
| Māori names used in this work | Scientific name and locality |
| Kapokapowai North (KN) | *Uropetala carovei*, Te Ika-a-Māui |
| Kapokapowai South (KS) | *Uropetala carovei*, Te Waipounamu (below 1.2 km) |
| Kapokapowai Montane (KM) | *Uropetala chiltoni*, Kā Tiritiri-o-te-Moana |

Despite its young geologic age, Te Moana o Raukawa has been proposed as a driver of allopatric speciation and population divergence in several taxa including amphipods (Stevens & Hogg, 2004), Weka (terrestrial birds) and their ectoparasites (Trewick et al., 2017), and skink lizards (O’Neill et al., 2008). The tectonic uplift of the Kā Tiritiri-o-te-Moana mountain range likely opened niches and acted as a driver of diversification in flora (Thomas et al., 2023) and fauna (Buckley et al., 2024) of Te Waipounamu. Most lineages of alpine flora arrived in the later Tertiary period, and the environmental upheaval in the Pliocene and Pleistocene drove diversity in these lineages (Thomas et al., 2023). Most alpine invertebrate fauna are relatively young, with ages corresponding to the increased uplift of the Kā Tiritiri-o-te-Moana range 5 million years ago (Ma), and the development of a permanent snow line just over 1 Ma (Buckley et al., 2024). Similar to the flora, it is broadly understood that most invertebrate fauna evolved from lineages already on the island as niches opened with the increased elevation (Buckley et al., 2024), although it has been hypothesized that invertebrates have ridden tectonic uplift to higher altitudes (Heads, 2017).

### Kapokapowai as an Important Study System in Island Biogeography

Aquatic insects have been used to better understand the past climate and geology of Aotearoa, and are useful models in the study of evolution and ecology (Córdoba-Aguilar et al., 2022), but they have been relatively underrepresented in biogeographic and broader biodiversity work (Leschen et al., 2023) and knowledge about biogeography remains limited for many taxa. One such group of aquatic macroinvertebrates are the dragonflies first referred to as Kapokapowai (among other names) by Māori (Grant, 2014), and later, named *Uropetala* by scientists from colonial Great Britain and Belgium (Selys 1858).

Kapokapowai are a particularly promising system for studying the impact of tectonic uplift and oceanic barriers in speciation as a member of the so-called “relict” (Ware et al., 2014) dragonfly lineage Petaluridae, a member of the previously diverse superfamily Petalurida. Petaluridae originated in the mid Jurassic, as sister to the species rich Gomphidae family, but comprising only 11 currently described species (see Tolman et al. (2024) and Cairns et al. (2025) for discussion of the status of one species of the genus *Petalura*) it is among the most species poor lineages of the order Odonata (Tolman et al., 2024). Despite inhabiting fen habitats which should readily promote fossilization, there is only one crown fossil Petaluridae, which is dated to the mid Cretaceous and closely resembles the extant species found at the fossil locality (Tolman et al., 2024). Thus, there is no compelling paleontological evidence that the Petaluridae were previously diverse. Speciation in this family appears to largely be driven by vicariance, as diversification in Petaluridae is strongly correlated with continental drift, and the expansion and contraction of inland seas and land bridges (Ware et al., 2014; Tolman et al., 2024).

Kapokapowai likely diverged from the Australian petalurids during continental split between Zealandia and Gondwana, and persisted through the Oligocene drowning of ∼18 Ma (Cooper and Cooper 1997; Ware et al., 2014; Tolman et al., 2024). Within Kapokapowai there are two currently described species, *Uropetala carovei* (Selys, 1843) which is thought to inhabit lower elevations of both Te Ika-a-Māui and Te Waipounamu (North and South Islands, respectively), and *Uropetala chiltoni* (Tillyard, 1921), which inhabits higher elevations of Te Waipounamu (South Island). Kapokapowai is the only lineage within Petaluridae where speciation is thought to have occured within the past 5 Ma (Tolman et al., 2024), thus the implications of geographical changes on species and population diversity can be studied in finer detail using this species complex than with other lineages of Petaluridae. The study of Kapokapowai can provide important hypotheses for the study of other “relict” lineages on island systems, defined by McCulloch and Waters (2019) to refer to phylogenetically “old lineages” found on geologically “young islands.” Island relicts have been considered to be a high priority for conservation biology (Grandcolas et al. 2014) and for the broader study of extinction and speciation more generally (Grandcolas et al. 2014).

### Māori Knowledge of Kapokapowai

In systems such as Kapokapowai, Indigenous knowledge is an important source of ecological and evolutionary information (Jessen et al. 2022). According to Māori cosmology, dragonflies and other insects are considered the children of Tāne Māhuta (the god of forests, birds and insects). Dragonflies appear in whakaairo (traditional carvings which often adorn the walls of Māori meeting houses) and waiata (songs) of Māori (Indigenous Peoples of Aotearoa). Stories about dragonflies also feature in pūrakau (legends) belonging to specific iwi (tribes) and hapū (subtribes). All insects, including dragonflies, were created before humans in Māori genealogical histories and are therefore tuakana (roughly translated as an elder family member) and are respected accordingly. Kapokapowai or Kapowai are names commonly-used for the Petaluridae of Aotearoa, which loosely translates to water snatchers, though there are other recorded names including Tītīwaiora, Kihihiwaru and Uruururoroa (Grant, 2014). Thus, these historical and cultural accounts provide valuable information about indirect as well as direct spatiotemporal distribution records of Kapokapowai. As this work was conducted with the express permission of the iwi who own the lands of Aotearoa, we refer to these dragonflies by the Māori name, Kapokapowai, in our work here. We also note that the name Kapokapowai was used to describe these dragonflies in the literature by Taylor (1855), italicizing “*Kapokapowai*” as to explicitly refer to a genus, three years prior to the first use of *Uropetala* by Selys (Grant, 2014; Taylor, 1855, 1870), and Kapokapowai has continued to be used to refer to dragonflies in this genus in scientific literature (Durrant, 2024; Fong, 2024; E. R. Tolman et al., 2024). However, we are not making any change to the binary taxonomic nomenclature of this genus here. In referring to Kapokapowai by their Māori name in this work, we designate *U. chiltoni* as Kapokapowai Montaine (KM), and *U. carovei* from either island as Kapokapowai North (KN) and Kapokapowai South (KS; Table 1), representing the three putative species in the Kapokapowai species complex.

### The Kapokapowai Species Complex

Debate about whether this complex should appropriately be considered to have two species has been ongoing for decades (Andress, 1998; L. S. Wolfe, 1952; Winstanley & Rowe, 1980). According to Tillyard’s original description, KM is characterized by large pale blotches on its labrum, which are not present in KN and KS, and by black femur leg segments, which are brown in KN and KS. However, variation in the splotches of the labrum have been used to suggest that hybrid populations between KM and KS can be found at the north end of Te Waipounamu (Winstanley & Rowe, 1980). Behavioral work suggests that female KM lay all their eggs in one plant, and in repeating this egg laying process females tend to badly damage their wings; such damage has not been seen in KN or KS (W. J. Winstanley, 1981). This behavioral work should be replicated to confirm that this is a species–wide behavior. It has also been suggested that KN and KS may be different species, as KN on Te Ika-a-Māui may have broader superior appendages and larger adult body size than their counterparts on Te Waipounamu (L. S. Wolfe, 1952). Two different karyotypes have been identified for this genus, but it is unclear if this is due to laboratory error (Kuznetsova & Golub, 2020); no other attempts have yet been made to determine the number of species in this genus using molecular data. Based on known morphological variation and behavioral data (L. S. Wolfe, 1952; Winstanley & Rowe, 1980), the dragonflies known as Kapokapowai are hypothesized here to comprise between one and three species, the diversification of which was shaped by both Te Moana o Raukawa (Cook Strait) and Kā Tiritiri-o-te-Moana (Southern Alps).

Here, we use population genomics, alongside scanning electron microscopy images of secondary male genitalia, and environmental niche models to determine the number of species and the population dynamics within Kapokapowai and to study how Kā Tiritiri-o-te-Moana and Te Moana o Raukawa may have driven speciation in this genus, and consider how our findings may inform evolution on Aotearoa more generally

## Materials and Methods

### Population genomics

#### Data acquisition

We utilized molecular data from a total of 37 dragonflies, 18 KM, 7 KN, 5 KS, and 7 *Phenes raptor* which were used as an outgroup (Supplemental table 1). We generated a 93 locus, 20kb (on average) nucleotide Anchored-Hybrid Enrichment (AHE) read library (Tolman et al., 2024) for 16 KM, and all other sequenced individuals, following the protocol used by Goodman et al (2023) and Tolman et al. (2024). Briefly, we extracted DNA from one leg of each specimen, which was removed using sterilized forceps, in the case of nymphs and adults. The entire thorax was used to extract DNA from exuvia. We incubated the tissue in proteinase K for one week at 55C, and then followed the manufacturers protocol for the remainder of the DNA extraction using the Zymo DNA Micro-prep kit (Hilden Germany). We sent the DNA extract to RAPID Genomics (Gainesville, Florida) for library preparation and sequencing. Additionally, we incorporated in the current study previously generated 1000 locus (which contained the 93 targeted loci for the 35 individuals sequenced here) AHE read libraries for 2 KM individuals, which were used in (Tolman et al., 2024) to estimate phylogenetic relationships of all species of Petaluridae.

#### Population Structure Analysis

We mapped the 37 sequenced AHE read libraries to a previously compiled soft-masked whole genome assembly (NCBI GenBank assembly GCA_036926305.1) of an individual KS (Tolman et al., 2023) with BWA2 (0.7.17-r1188; H. Li & Durbin, 2009) using default parameters, and calculated mapping statistics, average depth and converted the output to a sorted BAM file with samtools v1.16.1 (Danecek et al., 2021). We called variants with bcftools v1.15.1 (Danecek et al., 2021), using the options: `bcftools mpileup -a AD,DP,SP -Ou -f reference.fasta Input_bam | bcftools call - f GQ,GP -mO z -o output`. We then filtered the resulting VCF file for a minimum quality score of 30, and a minimum and maximum depth of 10 and 350 respectively to capture the region targeted with hybrid probes and the adjacent flanking region that is also sequenced and often more variable (Goodman et al., 2023).

To visualize population structure we performed a PCA analysis on a covariance matrix generated from the called SNPs with VCFtools v0.1.16 (Purcell et al., 2007) without pruning for linkage-disequilibrium (LD). To test if the outliers in the PCA could be a result of missing data, we identified the missingness of all individuals in the unpruned VCF file which excluded *Phenes*, and compared the missingness of KM with positive and negative PC1 scores. To model the ancestry of each sampled individual, we generated admixture plots for each putative species from admixture models assuming 2-7 ancestral populations with ADMIXTURE v1.3.0 (Skotte et al., 2013) and compared model fit using a cross-validation approach. The admixture graph under the best fitting models and the PCA plots were generated in R v2024.04.2+764 using mapmixture v1.1.0 library (Jenkins, 2024). Additionally, we calculated the F_ST_ between and within each of the three putative species both using VCFtools v0.1.16 (Purcell et al., 2007). To test for a correlation between genetic and geographic distance among our sampled individuals, we generated pairwise genetic distance matrices between sampled individuals with VCFtools v0.1.16 (Danecek et al., 2021), and then used the Mantel test in the vegan package in R v2024.04.2+764 (Oksanen et al., 2024), with 1,000 permutations to determine the significance of the correlation between geographic and genetic distance.

Genetic linkage can bias the recovery of population structure, as can inappropriately pruning sites (Bercovich et al. 2025). To determine the impact of genetic linkage on this dataset, and test the appropriateness of considering SNPs without LD pruning we pruned linked sites with PLINK v1.9.0-b.8 (Purcell et al., 2007) using the flags `--indep-pairwise 10 5 0` and re-ran the PCA analysis using the resulting VCF file.

#### Species delimitation

To delimit species based on our SNP data, we used the SPEEDEMON v.1.1.1 (Douglas & Bouckaert, 2022) pipeline in BEAST2 v2.7.7 (Bouckaert et al., 2014), using all 395 biallelic recovered SNPs after filtering.t,. We used snapper v1.1.4 (Stoltz et al., 2021) to run the SPEEDEMON pipeline on SNPs as opposed to alignments, and BICEPS v1.1.2 (Bouckaert, 2022) to implement a Yule Skyline Collapse Model to test for species delimitations, implemented with a chain length of 10^6 draws. We designated KM as a reference species, with KN and KS as putative species. We used the mean pairwise percent genetic divergence between individuals of KM (∼.54), as calculated from the VCF file using the pandas (McKinney, 2010) library for data manipulations and numpy (Harris et al., 2020) for numerical computations, as the prior mean on the theta prior (Douglas & Bouckaert, 2022). We used an alpha of 2.0, and beta of 0.26942 to achieve this mean in the Gamma distribution of the theta prior (see Stoltz et al., 2021 for a full description of these options), and followed the developer’s suggestions for all other parameters in the BEAST2 run (Stoltz et al., 2021). We independently ran the analysis two times to test for model convergence. We computed the posterior mean and upper credible (95%) limit of the Gelman-Rubin-Potential-Scale-Reduction Factor (PSRF) of each parameter, using the coda v0.19.4.1 library (Plummer et al. 2006), in r4.4.1(R Core Team 2021), to test for adquate model convergence. We also calculated the Effective Sample Size (ESS) for both runs in TRACER v1.7.2 (Rambaut et al., 2018) = to test for adequate mixing (at ESS >150) and model convergence (Rambaut et al., 2018).

#### Divergence time estimation

We utilized SNAPPER v1.1.4 (Stoltz et al., 2021) implemented in BEAST2 v2.7.7 (Bouckaert et al., 2014) to determine when the three putative species diverged, and to further explore which populations are most closely related. We used a different SNP set for this analysis, which included seven sampled *Phenes raptor* to use as an outgroup. The sequencing and variant call pipeline used to include these samples was identical to the protocol listed earlier. We used PLINK v1.9.0-b.8 (Purcell et al., 2007) with the flags `-- indep-pairwise 10 5 0.2` to prune linked sites from the VCF containing both *Phenes* and Kapokapowai. We used this pruned input file, with uniform age constraints of 113-135 Ma for the root age and of 0.1-15 Ma for the age of Kapokapowai (as previously identified by Tolman et al. 2024), to generate the snapper input file with a MCMC chain length of 10^7. SNAPPER can only implement a strict-clock model, a reasonable assumption for recently diverged populations and species, and (Stoltz et al., 2021), and we did not adjust the default tree model employed by SNAPPER. We calculated ESS for both runs in TRACER v1.7.2 (Rambaut et al., 2018) to test for adequate mixing, and computed the posterior mean and upper confidence (95%) limit of the PSRF of each parameter, using the coda v0.19.4.1 library (Plummer et al. 2006), in r4.4.1(R Core Team 2021), to test for adequate model convergence.

#### Tests of gene flow

As the resulting tree and species delimitation analyses supported KN and KS as sister, we tested for evidence of introgression between KM and KN and KM and KS with an ABBA-BABA test from Dsuite v0.5 r57, which is optimized to work on lineages with shorter evolutionary time scales. We used the unpruned VCF file which included *Phenes raptor* as input (Malinsky et al., 2021), and used *P. raptor* as an outgroup (E. R. Tolman et al., 2024).

### Morphological analysis

To identify potential morphological synapomorphies of the putative species of Kapokapowai we evaluated the male secondary genitalia. We specifically looked at the vesica spermalis, or sperm pump. This structure is located on the ventral side of the second abdominal segment in male Odonata, to which sperm is transferred prior to copulation, and may provide informative morphological characters for species determination (Kennedy, 1922; May, 1997; Ware, 2008). The vesica spermalis of Odonata is composed of four segments that are variable between species in their relative size and shape. In Petaluridae, a barb on the second segment near the junction with the third segment and the horns on the lateral side of the third segment have been used to support intrageneric difference (E. Tolman, 2024). We imaged the vesica spermalis (later referred to as penes) from two males from each putative species to try and identify more definitive diagnostic features for species delimitation. We followed the protocol for scanning electron microscopy outlined by Tolman et al. (2024). Briefly, penes were mounted laterally and dorsally on aluminum stubs and coated with gold palladium using a Cressington 108E sputter coater and specimens were then imaged using an S-7400 Hitachi scanning electron microscope.

### Environmental niche modeling

#### Occurrence Records

We acquired occurrence records of adult KN, KS, and KM from the Global Biodiversity Information Facility (GBIF)(available in Supplemental data). We selected occurrences from GBIF possessing preserved museum samples and research grade observations, which include verified latitude and longitude coordinates, a photograph of the sighting, observation date, and at least ⅔ agreement on species identification by the community. We filtered occurrences by removing sightings from erroneous localities (middle of the ocean, locations of large museums). We divided occurrences for KN and KS between Te Ika-a-Māui and Te Waipounamu (North and South Islands) to be used as individual models (Supplemental Table 3).

#### Environmental Data

We acquired averaged sets of environmental predictor variables for modeling consisting of purely bioclimatic variables. All analyses were conducted using the statistical programming language R v. 4.1.2. We acquired environmental rasters at 2.5 arc-second resolution (∼5km at the equator) from the CHELSA climate database v2.1 (Karger et al., 2017, 2023). We downloaded the ‘Anthropocene’ bioclimatic dataset (1979 – 2013) consisting of 19 bioclimatic variables which follow Worldclim v2 (Fick & Hijmans, 2017). We omitted bioclimatic variables 8 and 9 after visual inspection of interpolation discontinuities (Booth, 2022). Although such spatial artifacts were minor within our initial study extent, we observed major breaks of climate smoothing when we extrapolated our model to a larger spatial extent within the last glacial maximum and interglacial periods.

#### Model Construction

We spatially thinned occurrences to the resolution of our environmental variables to reduce the effects of sampling bias and artificial clustering (Veloz, 2009). We thinned occurrences by 5km to match the spatial resolution of our environmental data. We chose study extents for occurrences of KN and KS found in Te Ika-a-Māui and Te Waipounamu, defined as polygons around all localities between both islands respectively; we chose the same study extent for occurrences of KM found in Te Ika-a-Māui. We chose these study extents to include potential suitable habitat, while excluding large areas outside the species’ dispersal limitations (Peterson & Lieberman, 2012). Within this extent, we randomly sampled 50,000 background points for modeling and extracted their environmental values. We used these values to calculate correlations between variables using the ‘vifcor’ and ‘vifstep’ functions in the usdm package (Naimi, 2023) and filtered out variables with correlation coefficients higher than 0.7 and a Variance Inflation Factor (VIF) threshold of 10, both standard cutoffs in such analyses (Goodman et al., 2024).

To model the distribution of putative Kapokapowai species, we used the presence-background algorithm MaxEnt v3.4.4 (Phillips et al., 2017), using the R Package ENMeval 2.0.0 (Kass et al., 2021) for model building, parameterization, evaluation with different complexity settings, and reporting of results. We partitioned our data using the ‘checkerboard2’ strategy. To prevent our model from extrapolating beyond the bounds of training data, we clamped our models by omitting ranges of environmental data which fall outside of the training data. All final models were fitted to the full dataset. We tuned model complexity to find optimal settings using different feature classes, including linear (L), quadratic (Q), and hinge (H) as well as regularization multipliers 1 through 5 (higher numbers penalize complexity more) (Radosavljevic & Anderson, 2014; Warren & Seifert, 2011).

We assessed our model using averages of threshold-dependent (omission rate) and threshold-independent (Area Under the Receiver Operating Characteristic or AUC) discrimination performance metrics calculated on withheld validation data (Warren & Seifert, 2011). The 10th-percentile omission rate sets a threshold as the lowest suitability value for occurrences after removing the lowest 10% suitability values (Kass et al., 2021; Radosavljevic & Anderson, 2014). Validation AUC is a measure of discrimination accuracy that can be used to make relative comparisons between ENMs with different settings fit on the same data (Lobo et al., 2008; Radosavljevic & Anderson, 2014). Finally, we checked the results against the Akaike Information Criterion with correction for small sample sizes (AICc; calculated on the full dataset) (Warren & Seifert, 2011). To investigate model behavior, we examined marginal response to suitability. Marginal response curves show the modeled relationship of each variable individually with the occurrence data when all other variables are held constant (Phillips et al., 2017).

We made habitat suitability predictions for KM, and KN and KS from both islands using our ‘Anthropocene’ environmental predictor variables. We projected our modern-day models to a new study extent to encapsulate the potential habitat of Kapokapowai within the historic past, defined as a polygon around both Islands of Aotearoa. We generated thresholdless predictions by converting raw predictions to a scale of 0-1 to approximate probability of occurrence using the ‘cloglog’ transformation (Phillips et al. 2017). We also generated a threshold prediction, calculated from the 10-percentile omission rate from our model evaluation. To estimate habitat suitability of Kapokapowai within the geologic past, we generated predictions using environmental predictor variables from the PaleoClim dataset (Brown et al., 2018). We generated predictions within three distinct time periods: Mid-Holocene (8.3 – 4.2ka), Last Glacial Maximum (21ka), and the Last Interglacial (130ka).

#### Ordination Analysis

To determine niche differentiation between putative species, we compared niche overlap in occurrences of KN and KS from Te Ika-a-Māui and Te Waipounamu using an ordination framework by first reducing dimensionality within the datasets via a Principal Component analysis (PCA). Using the ‘espace_pca’ function in the *Wallace* v2.0.5 and *ade4* package v1.7 in R (Dray & Dufour, 2007; Kass et al., 2018)), we generated a Principal Components Analysis (PCA) using environmental variables from Kapokapowai occurrences and plotted with correlation loadings to infer the degree of influence particular environmental variables possess in the distribution of the 50,000 background points within niche space; we chose bioclimatic variables which were uncorrelated and shared among both Kapokapowai MaxEnt models. Using the ‘espace_occDens’ function in the package *ecospat* v3.2 (Di Cola et al., 2017, p. 2) an occurrence density grid was estimated for both the environmental values at each occurrence point and background extent points using a kernel density estimation approach. Niche overlap between occurrence density grids of environmental values at each occurrence point and background points were compared using Schoener’s D (Schoener, 1968) ‘espace_nicheOv’ function. Finally, using the ‘ecospat.plot.overlap.test’ function, we conducted a niche similarity test in which Kapokapowai niches from both islands are randomly shifted in the background extent and permuted 1000 times, and the niche overlap between the two are recalculated. A p-value <0.05 indicates that the niches from both time periods are significantly similar with each other.

#### Ancestral State Reconstruction of Elevation Ranges in Petaluridae

Having considered the niche space and population structure of low and high elevation Kapokapowai, then determined whether inhabiting mountains (KM) or lowlands (KN and KS) is the ancestral state for Kapokapowai–an important consideration in how these dragonflies are impacted by geography–by conducting an ancestral state reconstruction of the maximum elevation range for all of Petaluridae. We downloaded all occurrences of Petaluridae from the Global Biodiversity Information Facility (GBIF; GBIF.org, 2025) and determined the elevation of each occurence point using the elevatr v0.99.0 R package (Hollister et al. 2023). We determined the 90th percentile of elevation for each species of Petaluridae to minimize the impact of outlier observations, and classified the species as occupying high (>900 M), medium (400-900 M) or low (<400 M) elevations based upon this statistic. To model ancestral states we fit three variants of the Markov k-state model: Equal-Rates (ER), a single transition rate between all pairs of states, Symmetric-Rates (SR), distinct forward and backwards rates that are equal in reciprocal pairs; All-Rates-Differed (ARD), allowing each transition a unique rate. We fit each model with phytools v2.4-4 (Revell 2012) with estimated stationary frequencies. We also fit a custom model with equal rates between low and medium elevation, medium and high elevation, and a different rate between low and high elevation. We then compared model fit for the four models using AIC and used model averaging from ANCR to integrate node-state estimates across models according to their AIC rates. We used the dated species tree for Petaluridae (including the two currently described species of Kapokapowai) from Tolman et al. (2024) and added *Petalura pulcherrima* as sister to *P. ingentissima* at 14 Ma (within the confidence interval identified by Ware et al. (2014)). We tested the model with and without the fossil *Argentinopetala archangelskyi* which has been identified as sister to *P. raptor* (Tolman et al., 2024). To determine the elevation of this fossil at ∼115 Ma we converted the modern coordinates into the paleo-coordinates at 115 Ma using the palaeorate function in palaeoverse v1.4 (Jones et al. 2023) with grid rotation under the MERDITH2021 plate model. We used raster v3.6.31 (Hijmans, 2025) to load the 1° resolution Early Cretaceous Digital Elevation Model (DEM; Scotese and Wright 2018) and extract the paleo-elevation at the rotated point. As all states were equally probable at each internal node with and without *A. archangelskyi*, we reconducted searches with the averaged model using stochastic character mapping. To infer the frequency of changes between character states, we simulated 1000 discrete character histories under the selected averaged model, with empirical root-state frequencies set using the fitzjohn option in phytools v2.4-4 (Revell 2012). All aforementioned ancestral state reconstruction analyses were conducted in R v4.4.1 (R Core Team 2021).

## Results

### Population genomics

#### Population Structure Analysis

Mean sequencing depth varied from 0.02x to 0.19x, and missingness (calculated from the LD pruned VCF file) ranged from 0.01 to 0.27 (Supplemental Table 1). After filtering for mapping quality (> q30) and depth (10-350), we retained 365 SNPs for the initial population structure analysis. The weighted F_ST_ values were estimated at 0.23 between KM and KS, 0.29 between KM and KN and 0.11 between KN and KS (Table 2). We then separated the data into subsets as follows by their distributions along PC1 (Fig 1A): KM (all individuals), KM (negative PC axes), KN (Te Ika-a-Māui) and KS (Te Waipounamu). The F_ST_ values differed when only a subset of KM was used (Supplemental Table 3). There was no evidence that geographic separation was correlated with pairwise genetic distance within Kapokapowai (Pearson’s correlation coefficient *r* = 0.02462, p-value = 0.322).

**Figure 1:**
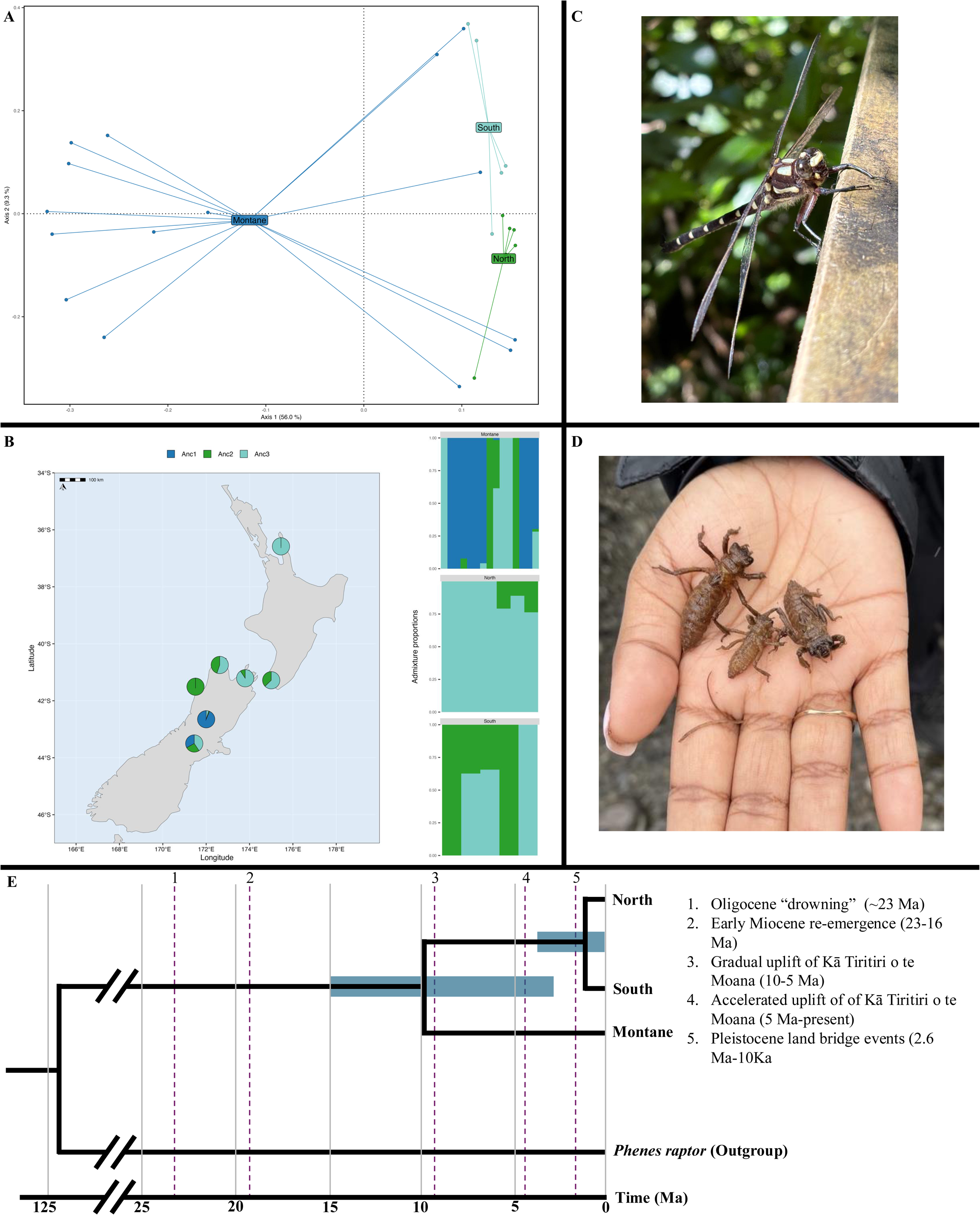
Population Genomics of Kapokapowai (A) PCA of Kapokapowai from SNPs, colored by putative species: Kapokapowai Montane (Blue), KN (Green) and KS (Teal). Each dot represents one individual with the distance from each point to the centroid of each putative species shown with a line. (B) Admixture analysis of Kapokapowai assuming three ancestral populations, with the ancestral populations 1-3 colored as blue, green, and teal respectively per each sampled individual (x-axis). Pie charts are mapped according to their proximate locality, and represent the proportion of each ancestral population in the gene pool of each locality. Ancestral population 1 is only found in KM. (C) Adult Female Kapokapowai North from Te Ika-a-Māui. (D) Nymphs of Kapokapowai Montane. (E) Dated tree displaying the divergence between the three putative species and the outgroup *Phenes raptor*. Notable geographic events in Aotearoa are annotated on the tree, including the Oligocene drowning when Aotearoa was reduced to 18% of its current size (Cooper and Cooper 1997), the reemergence of taxa following the drowning, the uplift of Kā Tiritiri o te Moana, and the Pleistocene land bridge events between Te Ika-a-Māui and Te Waipounamu.

**Table 2:** *F_st_ values between putative species of Kapokapowai*, using 365 called SNPs. Kapokapowai North (KN), Kapokapowai Montaine (KM), and Kapokapowai South (KS) are abbreviated. The results show stronger segmentation between KM and KN and KS.

| Pairwise comparison | Mean $F_{ST}$ | Weighted $F_{ST}$ |
| --- | --- | --- |
| KM-KS | 0.20 | 0.23 |
| KM-KN | 0.27 | 0.29 |
| KN-KS | 0.05 | 0.11 |

**Table 3:** gene flow between Populations of Kapokapowai.

| P1 | P2 | P3 | Dstatistic | Z-score | p-value | f4-ratio | BBAA | ABBA | BABA |
| --- | --- | --- | --- | --- | --- | --- | --- | --- | --- |
| KN | KS | KM | 0.15 | 3.74 | < 0.01 | 0.08 | 25.89 | 28.52 | 20.86 |

The centroid of KM was distal from the centroids of KN and KS across PC1, while the centroid of KN and KS differed along PC2 (Fig. 1A). Despite the differing centroids across PC1 and PC2, there was overlap in the space occupied between all three putative species (Fig. 1A). The missingness of KM individuals with a positive PC1 ranged from 0.02 to 0.17, while the missingness of KM individuals with a negative PC1 value ranged from 0.07 to 0.16 (Supplemental Table 3). 118 SNPs were retained after LD pruning the VCF, although the resulting population structure visualized through PCA (Supplemental Fig. 1) largely reflected the structure generated from pruned SNPs (Fig. 1A), with a slightly reduced distance between KN and KS.

Because of the low cross validation-error indicating comparable fit to other tested models (Supplemental table 5), and the three clusters identified in our PCA analysis, we provide the three ancestral populations model in our main figure and discussion (Fig. 1B), but also provide assumptions of K = 2, and K = 4 (Supplemental fig. 2). Ancestral population 1 was only present in KM, while ancestral populations 2 and 3 were present in all three putative species (Fig. 1B).

#### Species delimitation analysis

All parameters in the SPEEDEMON analysis showed adequate mixing (ESS > 400), and model convergence (PSRF = 1.0 with 95% CI <1.01 for all parameters). The species were recovered as currently described, specifically KM as *U. chiltoni* and KN and KS as *U. carovei*, with very high (99%) posterior support.

#### Divergence time estimation

Including the outgroup *Phenes raptor* we recovered 727 SNPs and retained 204 of these SNPs after LD pruning. The SPEEDEMON runs showed adequate mixing (ESS>600 for posterior, likelihood, prior, lambda and tree height; ESS >150 for clock rate) and strong evidence of convergence (PSRF=1.0 with 95% CI <1.06 for all parameters). In the dated tree Kapokapowai and *Phenes* shared a Most Recent Common Ancestor (MRCA) between 113.0 and 133.7 Ma, KM shared a MRCA with KN and KS between 2.6 and 15.0 Ma, while KN and KS diverged between 0.1 and 3.2 Ma (Fig. 1E; Supplemental Fig. 2).

#### Tests of gene flow

The ABBA-BABA test, implemented in D-suite using all 727 SNPs identifie devidence of gene flow between KM and KS (p < .01; Table 2)). It is important to note that this test cannot determine the directionality of gene flow.

#### Morphological analysis

The “horns” of KM (see figure 2, feature 1) are downcurved at the tips in a way not observed in any of the three KN or KS individuals (Fig. 3). The angle and curvature of the barb (figure 2, feature 2), and the presence of a “knob” near it (figure 2, feature 3) differs between individuals, but does not appear to be fixed between any populations. One individual of KN had a knob near the barb.

**Figure 2:**
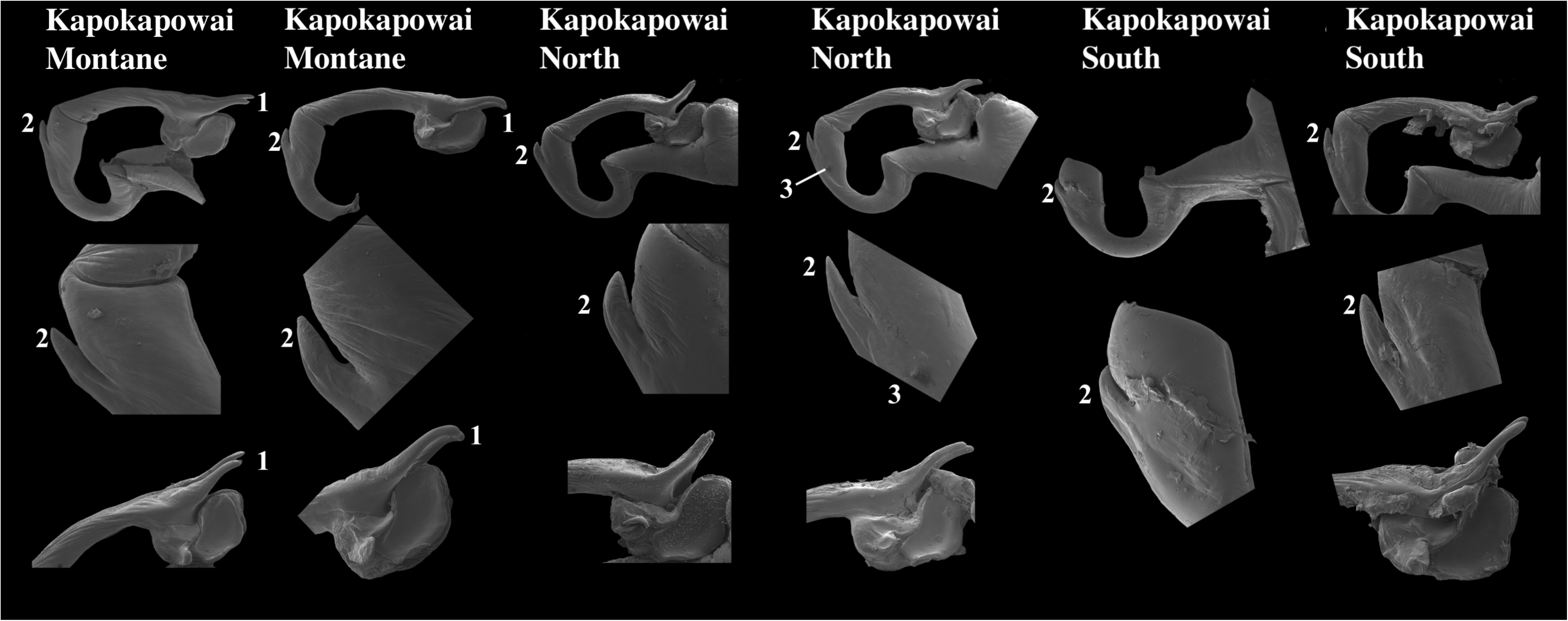
Male Secondary Genitalia of the Three Hypothesized Kapokapowai Species Scanning electron microscopy. Images of male penes from six sampled Kapokapowai (two from each putative species). Divergent features include (1) curves at the end of the horns only found in Kapokapowai Montane, (2) the barb near the joint of the second and third segments, and (3) a knob found near the barb from one individual from Kapokapowai North.

**Figure 3:**
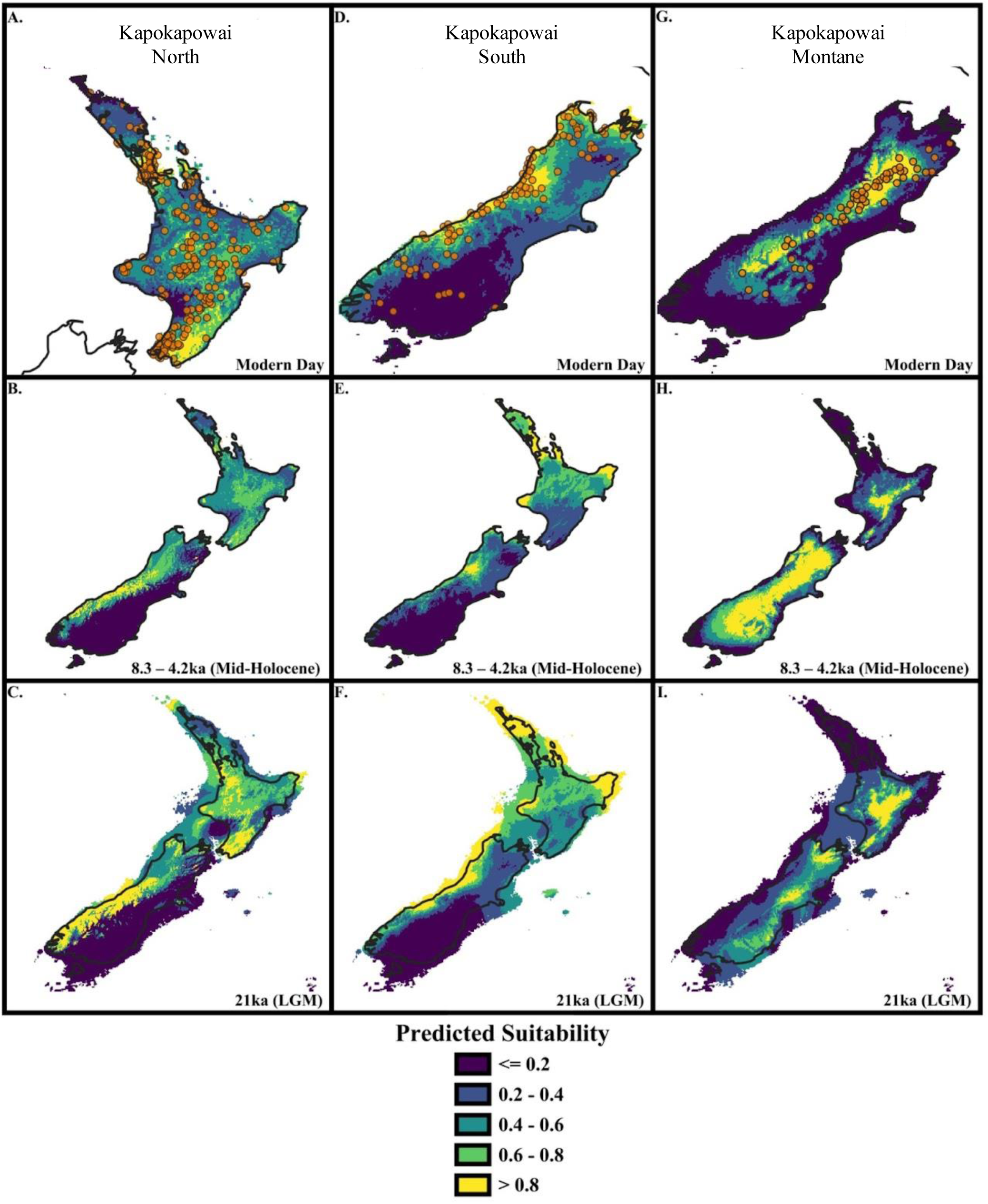
Niche Models of Kapokapowai Populations. MaxEnt predictions for Kapokapowai projected to the modern day using our ‘Anthropocene variables’, mid-Holocene (Northgrippian: 4.2 – 8.326ka) generated from the PaleoClim dataset (Brown et al. 2018), and Last Glacial Maximum (LGM)(21ka) and Last Interglacial (LIG)(130ka) generated from the CHELSA database (Karger et al. 2017). Predictions were derived from the optimal model using the criterion of mean valuation AUC (AUC_val_) values being the highest, and 10% omission rate being the lowest. Predictions were transformed using the ‘cloglog’ function in Maxent v3.4.4 (Phillips et al., 2017), in which raw values are converted to a range of 0 - 1 to approximate a probability of occurrence. Brighter colors (yellow, green, blue) indicate areas of higher suitability (higher probability of occurrence), while darker colors (violet) indicate areas of lower suitability (lower probability of occurrence).

#### Ecological Niche Modeling

Optimal model settings varied considerably for Kapokapowai based on 10-percentile omission rate and mean validation AUC metrics. Based on the results of the collinearity analysis, we used the following six predictor variables to build all models: Isothermality (bio03), temperature seasonality (bio04), maximum temperature of the warmest month (bio05), precipitation seasonality (bio15), precipitation of the warmest quarter (bio18), and precipitation of the coldest quarter (bio19). Models varied in complexity with expressing either hinge-loss, or linear-quadratic feature classes with regularization multipliers of 3-4 (higher complexity penalty). Mean validation AUC was adequate for KS and KM (0.75 and 0.86 respectively), while 10% omission rate was low relative to its expected value of 0.1 (0.11 and 0.09 respectively) (Supplemental Table 2). Furthermore, optimal models for KS and KM possessed low amounts of non-zero lambdas (coefficients) and AICc values suggesting low amounts of model overinflation (Supplemental Table 2). Optimal models for KN performed poorly, possessing an unacceptably low AUC value (0.44) and high 10% omission rate (0.16) (Supplemental Table 2).

Optimal models retained 4 – 6 predictor variables after regularization, and relied mainly on contributions from Isothermality (bio03), temperature seasonality (bio04), maximum temperature of the warmest month (bio05), and precipitation of the wettest and driest quarters (bio18 and bio19 respectively) (See Supplemental Fig. 3). Response curves of climate variables revealed differing relationships with suitability among Kapokapowai models (Supplemental Fig. 1). Response curves for KN expressed jagged optimal ranges, with habitat suitability not following a smooth or continuous gadient. Temperature Seasonality (bio03) and precipitation of the coldest quarter expressed positive linear/hinge relationships with suitability among all three Kapokapowai models, while precipitation of the wettest quarter (bio18) expressed negative linear/hinge relationships with suitability for KN and KS; precipitation of the wettest quarter possessed no relationship in KM. Isothermality (bio03), and maximum temperature of the warmest month (bio05) expressed opposite relationships with suitability within Kapokapowai. Isothermality (bio03) expressed positive linear or linear/quadratic relationships with suitability within KM and KN, but a negative linear relationship with KS. Maximum temperature of the warmest month (bio05) expressed negative linear or quadratic/linear relationships with suitability within KM and KS but a positive linear relationship within Kapokaowai South (Supplemental Fig. 1).

Predictions using our ‘Anthropocene’ environmental variables for KS within Te Waipounamu revealed areas of high suitability along the coast (Westland, Nelson and Marlborough provinces) (Fig 3D). Suitability was highest for KM across the Southern Alps (interior Nelson and Marlborough provinces) (Fig. 3A). Predictions for Kapokaowai North within Te Ika-a-Māui revealed areas of high suitability within the interior (Wellington and Hawke’s Bay, along the Tararura, Ruahine, and Kaimanawa Mountain Ranges), as well as the Northern portion of the island within Auckland (Fig 3G). Threshold predictions reflect our ‘cloglog’ predictions (See Supplemental Fig. 3).

Predictions generated from our paleoclimate variables demonstrated variability in suitability over time for Kapokapowai compared to our modern-day ‘Anthropocene’ variables. For KM, suitability expands within the mid-Holocene (8.3 – 4.2ka), encompassing most of Te Waipounamu except for the coastal margins (Fig 3H.). Projection to the Te Ika-a-Māui within the mid-Holocene indicates suitable habitat within the island’s center (Te Ika-a-Māui Volcanic Plateau, Tararua Range, and Raukumara Range) (Fig 3H.). Within the LGM, suitability contracts within Te Waipounamu, with suitable habitat only being located within a small portion of the Kā Tiritiri-o-te-Moana (Southern Alps) and Kaikōura Mountain Ranges (Fig 3I.). Within Te Ika-a-Māui, suitability expands, encompassing most of the western and southern mountain ranges. Although a land bridge connected both Islands during the LGM (Te Moana o Raukawa), suitability was low (Fig 3I.) while our threshold predictions suggest suitable habitat across the strait (See Supplemental Fig. 4).

For KS within Te Waipounamu, suitability contracts within the mid-Holocene, being restricted to a small portion of the coast (northern Westland) (Fig 3E.). When projected to Te Ika-a-Māui, suitability is high within the Northern Cape and southwestern cape (Taranaki Province), and Auckland (Fig 3E). Within the LGM, suitability expands encompassing the northern coastal regions of Te Waipounamu (Westland and Nelson), as well as the Northern Cape, Taranaki Province, and Auckland within Te Ika-a-Māui (Fig 3F.). Suitability across the Te Moana o Raukawa is intermediate, with a few pockets of high suitability (Fig 3F.) while our threshold predictions suggest suitable habitat across the strait (See Supplemental Fig. 4).

For KN within Te Ika-a-Māui, suitability contracts within the mid-Holocene, with only a few pockets of high suitability within the southern coasts (Tararua Ranges), and Auckland (Fig. 3B.). When projected to Te Waipounamu, suitability is highest along the coastal ranges of southern Westland and Otago. Within the LGM, suitability expands to include most of the southern Mountain Ranges within Te Ika-a-Māui, as well as Kaimanawa Mountain Ranges within the middle of the Island (Fig 3C). Within Te Waipounamu, suitability slightly expands southward into the coastal ranges of Otago (Fig 3D). Suitability within the Te Moana o Raukawa remains intermediate with a small pocket of high suitability off the coast of Taranaki (Fig 3D.) while our threshold predictions suggest suitable habitat across the strait (See Supplemental Fig. 4).

#### Ordination Analysis

Environmental variables for KN and KSwithin Te Ika-a-Māui and Te Waipounamu expressed an intermediate degree of overlap in ordination analysis, but were statistically more similar than random (Fig. 4). Within our PCA analysis, the first principal component explained ∼34% of the variance with PC2 explaining ∼32%. Isothermality (bio03), temperature seasonality (bio04), and precipitation of the coldest quarter (bio19) loaded strongly on PC1, while precipitation of the warmest and coldest quarters (bio18 and bio19) loaded strongly on PC2 (Table 4). Environmental niche overlap for Kapokapowai was intermediate between Te Ika-a-Māui and Te Waipounamu (Schoener’s D = 0.24), with niche similarity tests showing observed overlaps being higher than 95% of simulated overlaps (*P* = 0.01), supporting niche similarity between both islands (Fig. 4).

**Figure 4:**
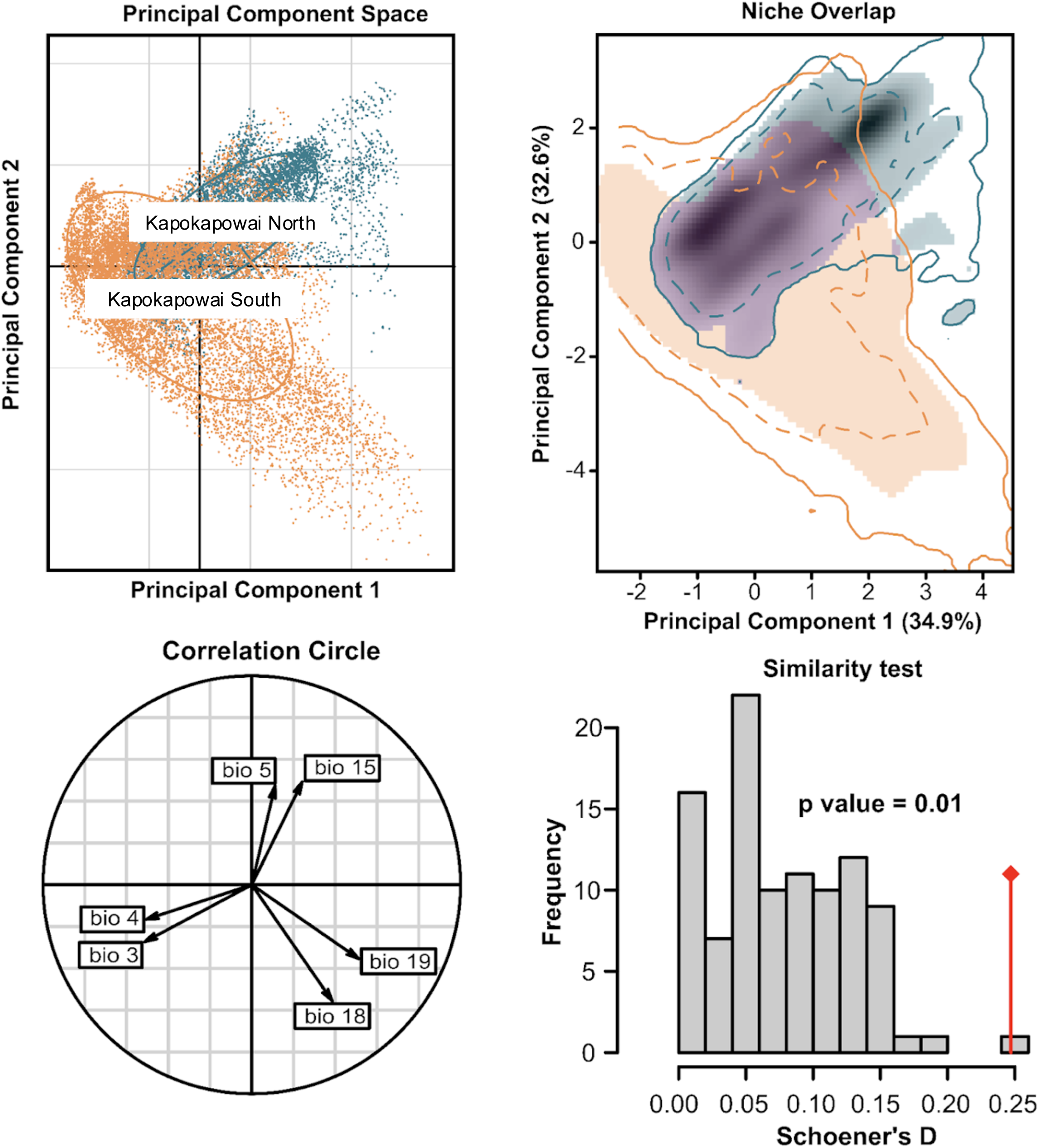
Niche Overlap of Kapokapowai North and Kapokapowai South. Ordination plot (PCA), niche overlap, correlation circle, and niche similarity tests showing niche differences between Kapokapowai North and South from Te Ika-a-Māui (North) and Te Waipounamu (South). Ordination plot represents principal component points of occurrence and background environmental values using type 2 scaling (Mahalanobis distance). PCA points for Te Waipounamu are in blue, while Te Ika-a-Māui is red. Solid contour lines in the niche overlap illustrate full range (100%) of climate space (‘fundamental niche’) while dashed lines indicate 50% confidence intervals. Contour lines of Te Waipounamu are in orange, and Te Ika-a-Māui is blue. Shading shows the density of species occurrences per grid cell of kernel density analysis (‘realized niche’), and violet pixels show niche stability (climate conditions occupied in both time periods). Orange shading indicates climate conditions only occupied by Te Waipounamu, while blue indicates climate conditions only occupied by Te Ika-a-Māui. Correlation circle indicates climactic variable loadings on the PCA space. Length and direction of arrows indicates influence and distribution of variables within PCA space. Bioclimatic variables which follow Worldclim v2 (Fick and Hijmans 2017). Histograms represent observed (red bar) and randomly simulated overlaps of niche similarity. The observed overlap is higher than 95% (p < 0.05) for both analyses supporting the niche similarity across both islands.

**Table 4:**
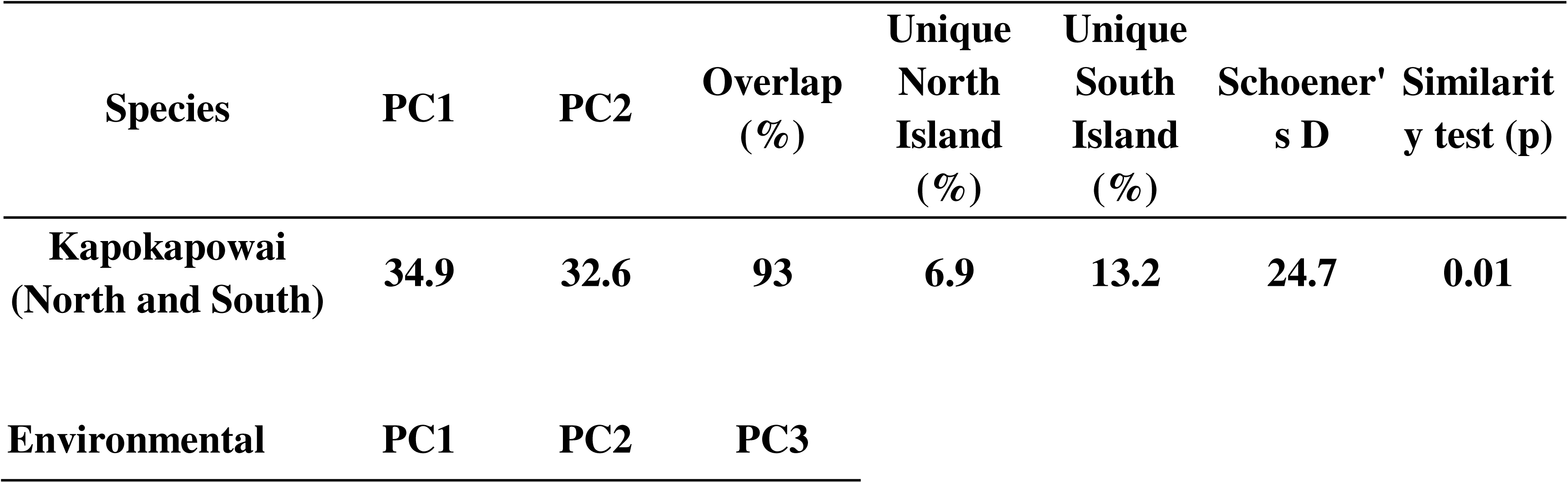
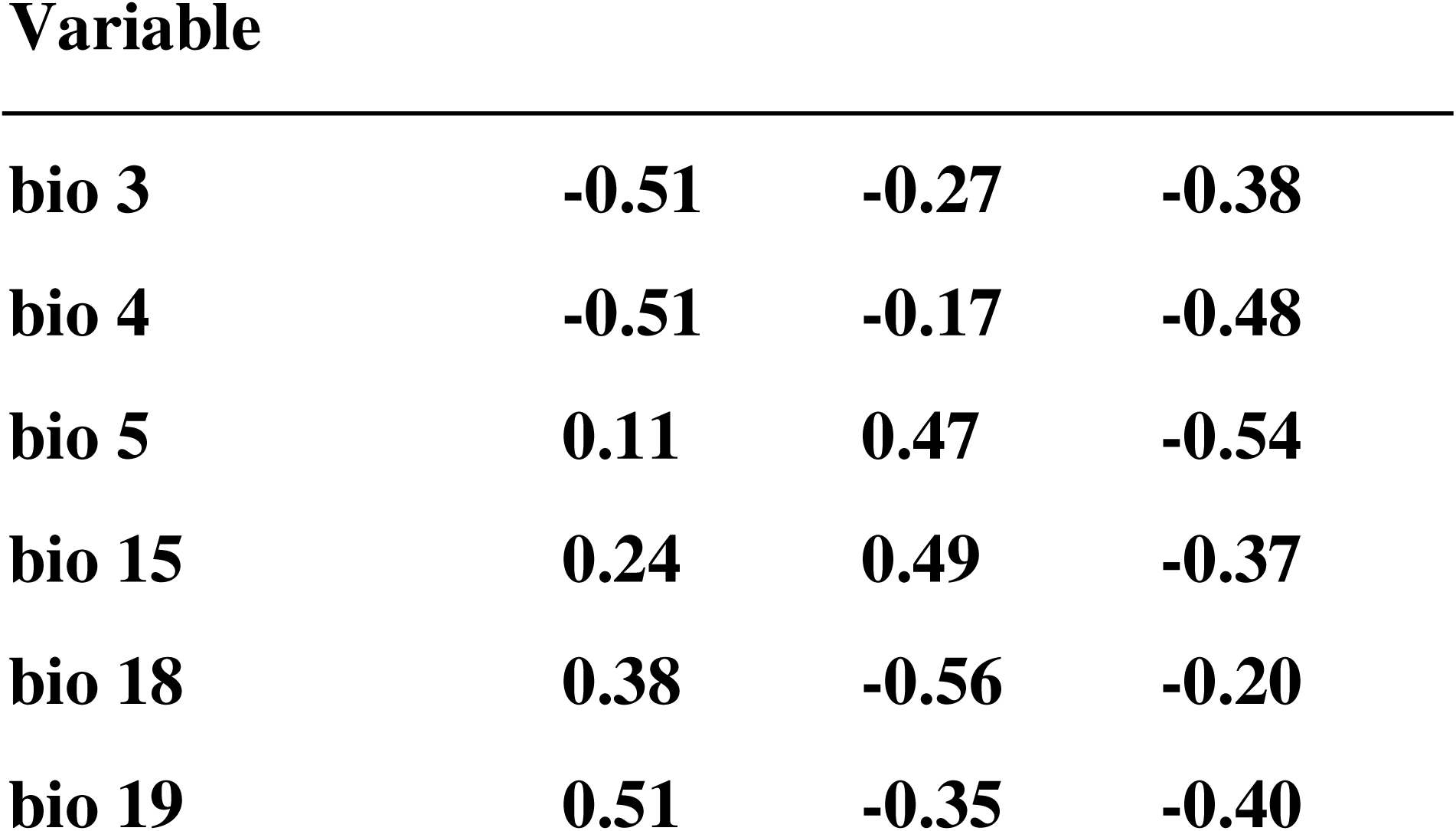
PCA loadings, importance of variables, percent overlap, percent of unique niche overlap from both islands, environmental similarity (Schoener’s D), and results of similarity tests from our EcoSpat analyses.

| Species | PC1 | PC2 | Overlap (%) | Unique North Island (%) | Unique South Island (%) | Schoener's D | Similarity test (p) |
| --- | --- | --- | --- | --- | --- | --- | --- |
| Kapokapowai (North and South) | 34.9 | 32.6 | 93 | 6.9 | 13.2 | 24.7 | 0.01 |
| Environmental | PC1 | PC2 | PC3 |  |  |  |  |

**Variable**
|  |  |  |  |
| --- | --- | --- | --- |
| **bio 3** | **-0.51** | **-0.27** | **-0.38** |
| **bio 4** | **-0.51** | **-0.17** | **-0.48** |
| **bio 5** | **0.11** | **0.47** | **-0.54** |
| **bio 15** | **0.24** | **0.49** | **-0.37** |
| **bio 18** | **0.38** | **-0.56** | **-0.20** |
| **bio 19** | **0.51** | **-0.35** | **-0.40** |

#### Ancestral State Reconstruction

The ER model was strongly supported by AIC (Supplemental Table 6) both with and without *A. archangelskyi*, indicating gains and losses of different elevation habitats are equally probable across the tree. All nodes were equivalent with near equal support for each elevational maxim, including the ancestral state of Petaluridae, in both maximum likelihood models and using stochastic character mapping, both with and without *A. archangelskyi* (Fig. 5). Across 1000 stochastic maps under the averaged model with *A. archangeslskyi* included in the tree, we inferred on average 235 instances of medium to high elevation and high to medium elevation with transition and 236 medium to low and low to medium transitions, meaning each of these transitions are expected to occur once every 2.9 million years for a given branch of Petaluridae on average. We inferred 49 high to low and low to high elevation transitions each, averaging one of each transition every 14.5 million years for a given branch of Petaluridae on average. The expected frequency of transitions per million years for a given branch of Petaluridae did not change with *A. archangelskyi* removed.

**Figure 5:**
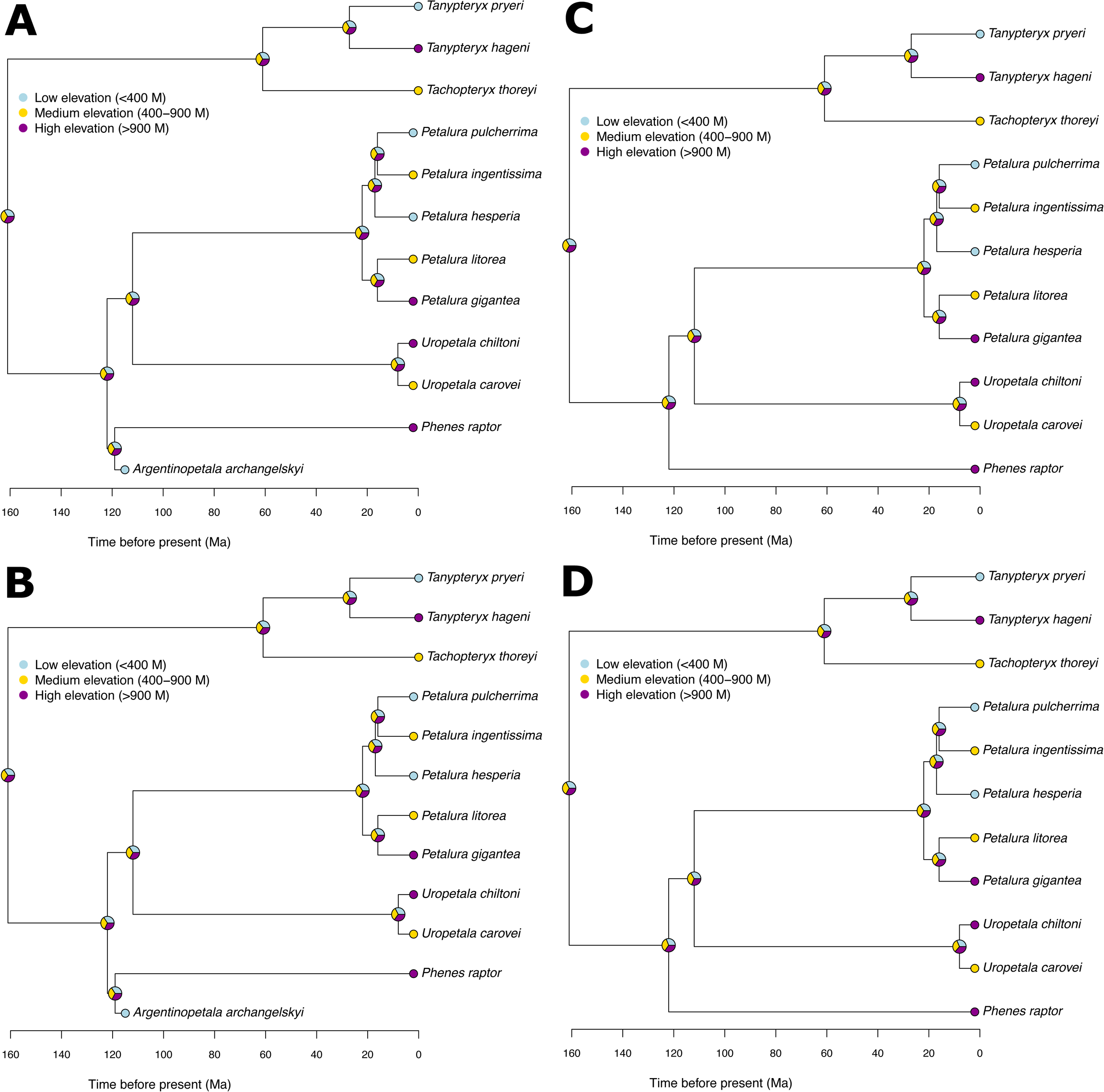
Ancestral State Reconstruction of the Maximum Elevation inhabited by Petaluridae. Maximum likelihood (ML) reconstructions and stochastic character mapping (SM) are shown both with (ML A, SM B) and without (ML C, SM D) the fossil taxa *Argentinopetala archangelskyi*, which is considered to be sister to *P. raptor* based upon wing characteristics (Tolman et al., 2024). In all models, all three ancestral states (high, medium, and low maximum elevation) are equally likely as the ancestral state at each node, including at the root.

## Discussion

### Historical Island Geography Underlies a Species Complex

As hypothesized, our results demonstrate that uplift and oceanic barriers have shaped the evolution of Kapokapowai. The divergence time analysis suggested that the MRCA (2.6-15 Ma) of all sampled Kapokapowai could very well have coincided with the accelerated tectonic uplift which occurred ∼5Ma, or the more gradual uplift from 10-5Ma (Fig. 1E). Intriguingly, a strong bottleneck in the ancestral lineage of KS was identified between 2.5-10Ma from a reference genome (Tolman et al., 2023) using the Pairwise Sequential Coalescent (PSMC; Liu & Hansen, 2017). This bottleneck could plausibly be reflective of speciation between KM and KN+KS. KM has higher genetic diversity than either KN or KS (Fig. 1A,B) supporting the hypothesis that this bottleneck was caused by a founder effect similar to many other systems (Anopheles gambiae 1000 Genomes Consortium et al., 2017; Auton et al., 2015; B. Li et al., 2024).

The genetic differentiation between KN and KS is much smaller than the differentiation between either population and KM (Table 2). Although Te Moana o Raukawa (Cook Strait) is currently a barrier to the dispersal of Kapokapowai, our species distribution models suggest that this has not always been the case (Fig. 4, Supplemental Fig. 4). Perhaps Te Moana o Raukawa has not been a barrier to the dispersal of Kapokapowai for a sufficient period of time for a strong genetic structure to evolve. On the other hand, the peaks of Kā Tiritiri-o-te-Moana have not contained suitable habitat for KN or KS in any of the tested time periods, and have likely served as a longer barrier to the dispersal of these populations, shaping the genetic structure currently observed in Kapokapowai. The physical boundary of the mountains is further compounded by the fact that the alpine habitat has shortened the flight season of KM, compared to KN and KS (L. S. Wolfe, 1952).

The data we have here support a hypothesis in which all sampled Kapokapowai are descendents of a population that colonized alpine habitats (the most supported pattern in the fauna of Aotearoa (Buckley et al., 2024), although there is speculation that some lineages may have ridden tectonic uplift to higher elevations (Heads, 2017)), while KN and KS descend from populations that later moved into lowland habitats. Our ancestral state reconstruction analysis of Petaluridae would support the former, as across 1000 simulations, transitions from high elevation to low and medium elevational maxims occurred on average every 2.9 million years on each branch. It is entirely plausible that such flexibility in adjusting to different elevations is a common trait in lineages that have persisted on Zealandia for tens of millions of years.This hypothesis of recolonization of the lowlands is not without precedent, as plant lineages have also been hypothesized to have colonized mountains, and then recolonized the lowlands of Aoteora (Thomas et al., 2023). A demographic model for KM from a reference genome, and revisiting the specimens sampled here with genome-wide resequencing data could be used to generate a demographic model for all populations of Kapokapowai, and a historical demographic model of dispersal and gene flow between populations using the site-frequency-spectrum (Noskova et al., 2023), both of which could be used to robustly test this hypothesis.

### Gene Flow in the Presence of Geographic Barriers and Implications for Species Delimitation

#### Evidence of Current gene flow

Despite the aforementioned geographic barriers, the PCA (Fig. 1A), admixture (Fig. 1B), and introgression analyses all suggest there is some level of interbreeding and movement between populations. Neither barrier seems to be entirely restricting gene flow, as evidenced by lower fixation (F_ST_ < 0.30) than might be expected given > 5 million years of separation between KM and KN and KS (Fig. 1E), and evidence of introgression between KS and KM.

One individual KM notably inherited 75% of its ancestry from ancestral population one, which is unique to KM, and 25% of its ancestry from ancestral population three (Fig. 1B). Thus, this individual is likely the offspring of an F1-hybrid back-crossed with an individual with 100% of its ancestry from population one, suggesting the genetic populations can breed with one another, and gene flow may still occur in the present.

#### Implications for species delimitation

Given the dynamics of gene flow, current species delimitations in this complex are difficult to make. Given the genetic differentiation between KM and KN and KS (F_ST_ >.23), our identification of a diagnostic characteristic of the curvature on the horns of KM (Fig. 2), and the strong posterior support (>99%) of the currently described species (*U. carovei* comprised of KN and KS and *U. chiltoni* comprised of KM) in our species delimitation analysis, we suggest that current species delimitations remain unchanged. Although KN and KS do show some genetic differentiation, they do not show evidence of niche divergence, and we do not believe that they merit a new species description at this point in time.

### Generating Species Distribution Models for Island Species

Although the utilization of GBIF occurrences in conjunction with genomic analyses provide the opportunity for highly robust niche models which have the potential to match population admixture or structuring among populations of odonates, a consequence is the underfitting of model performance due to the abundance, range, and density of the species itself. ENM results for KN revealed low AUC values, which is a continuous metric that provides discriminative ability in false positives and false negatives (Phillips et al. 2017). Since KN is found throughout the entirety of Te Ika-a-Māui, our models were unable to discern gradients of high and low suitability, resulting in poor model output. This pattern is also observed in our AICc value (Supplementary Table S2). Our thinned dataset consisted of 239 occurrences, sufficient to generate a robust model; the widespread distribution throughout Te Ika-a-Māui contributed to high AICc scores.

ENMs in the context of island biogeography is an area of ongoing study, as model extrapolation becomes poor for species endemic to small or singular islands (Sutherland et al. 2021, Goodman et al. 2024), limiting the incorporation of novel climatic conditions that could improve model fit. Furthermore AUC values may be skewed if only a few climatic variables drive the model, leading to oversimplified relationships with suitability (Phillips et al. 2017), as evidenced by the number of nonzero lambda coefficients within our KN model (2)(See Supplementary Table S2).

Interestingly, our 10-percentile omission rate metrics were more robust (0.16). The 10-percentile omission rate metric excludes the lowest 10 percent of raw occurrence values, thus creating a threshold within suitability. Our lower omission rates suggest minimal outliers in our dataset, suggesting that our predictions for KN on Te Ika-a-Māui are not overly generalist.

An aspect to consider in paleo-ecological modeling (PaleoENM) is the use of multiple lines of evidence to assess model fidelity. Incorporating supplemental analyses such as response curves and ordination analyses provide clarity to model over- or under-extrapolation, which can be interpreted as either model error or accuracy. Response curves demonstrate if the full environmental range of the species has been sampled for each species or population, while ordination analyses allow multidimensional visualization of the niche outside of a spatial context. In the case of our data, response curves, albeit jagged, map the full relationship of each climatic variable within the model, while our ordination analysis demonstrates statistical differences among the KN and KS populations. Overall, although genetic divergence was low among KN and KS populations, the ecological and distributional differences among both populations are the result in differences in climatic conditions, and not a result of poor model performance because of inappropriate apriori grouping of populations based on their presence on Te Ika-a-Māui or Te Waipounamu.

## Conclusion

To look at species diversity and the potential for speciation in a “relict” group such as the petaltail dragonflies offers a unique perspective in evolutionary biology, as well as island ecology. Analyses suggest that this family is extremely old, even for an order as ancient as Odonata, with some extant species persisting as independent evolutionary lineages since the Cretaceous era (Kohli et al., 2021; Suvorov et al., 2022; E. R. Tolman et al., 2024). With their reliance on fen habitats, Petaluridae are also somewhat limited in their range, compared to more generalist dragonflies that use ponds or rivers to complete their life cycle. Despite this, our results demonstrate that speciation in Petaluridae is associated with shifts to habitats at different elevations, in both Kapokapowai and other groups of petaltails, which may be associated with their long persistence. While species of Kapokapowai have potentially been separated for significant periods of time, either across islands or in different habitats within an island, it may be that their adaptations to these specialized fen habitats has limited the genetic variability that would lead to speciation. This may help to explain why this group, despite its age, continues to be the smallest family within Anisoptera.

## Acknowledgements

We would like to thank Thomas Beatty and Mahealani Kaloi for their support of this project. We would also like to acknowledge Dr. Milen Marinov for his generosity in providing Odonata specimens, as well as the Department of Entomology at the Smithsonian Institution’s National Museum of Natural History, Florida State Collection of Arthropods, and American Museum of Natural History for providing specimens used in this analysis.

## Data availability

Filtered VCF files with and without *P. raptor*, and log and xml files from the BEAST2 runs have been uploaded to figshare (10.6084/m9.figshare.27896910). The Raw reads have been uploaded to the sequencing read archive (bioproject PRJNA1190089).

