## Supplementary Materials for "Elevational and Oceanic Barriers Shape the Distribution, Dispersal and Diversity of Aotearoa’s Kapokapowai (*Uropetala*) Dragonflies"

| Supplementary Table 1: Specimens Sequenced for Molecular Analyses | | | | | | | | | | | |
| --- | --- | --- | --- | --- | --- | --- | --- | --- | --- | --- | --- |
| Specimen type | Genus | Species | Population | Locality | Read Identifier | Average depth | SNPs missing-ness (LD pruned VCF) | Lat | Long | Collection Date | Collected by |
| Adult | *Phenes* | raptor | - | Malleco Prov.; Angol | WE07 | 0.05 | 0.04 | - | - | 10/19/65 | full names will be added on acceptance |
| Adult | *Phenes* | raptor | - | Nuble Prov.; Alto Tregualemu, ca 20 km SE Chovellen, 500 M | WE08 | 0.04 | 0.01 | - | - | 12/3/81 | full names will be added on acceptance |
| Adult | *Phenes* | raptor | - | Linares; Embalse Bullileo | WE09 | 0.06 | 0.04 | -36.18 | -71.25 | 1/12/94 | full names will be added on acceptance |
| Adult | *Phenes* | raptor | - | 20 km N. of Vallenar Atac. | WE10 | 0.04 | 0.06 | - | - | 10/22/82 | full names will be added on acceptance |
| Adult | *Phenes* | raptor | - | Cacapoal Prov; Rio Peuco, Pilay, ca. 45 km S Santiago, 800m | WF01 | 0.07 | 0.03 | - | - | 11/26/81 | full names will be added on acceptance |
| Adult | *Phenes* | raptor | - | Aysen Prov; Chaiten | WF02 | 0.05 | 0.04 | - | - | 12/1985 | full names will be added on acceptance |
| Adult | *Phenes* | raptor | - | Naheulbuta, Pichinahuel | WF03 | 0.10 | 0.03 | -37.48 | -73.02 | 12/19/93 | full names will be added on acceptance |
| Adult | *Uropetala* | carovei | KN | Coppermine Creek, Ruahini Forest Park | WF05 | 0.07 | 0.02 | -40.14 | 175.53 | 2/11/98 | full names will be added on acceptance |
| Adult | *Uropetala* | carovei | KN | Catchpool Stream Rimutaka Forest Park, S. Wainuiomata | WF06 | 0.05 | 0.03 | -41.21 | 174.55 | 2/18/01 | full names will be added on acceptance |
| Adult | *Uropetala* | carovei | KN | Near Top, Smiths Creek Track, Tararua Forest Park, 16km NE Upper Hutt | WF07 | 0.04 | 0.03 | -41.04 | 175.13 | 2/8/98 | full names will be added on acceptance |
| Adult | *Uropetala* | carovei | KN | Tokaanu Thermal Park | WF08 | 0.08 | 0.01 | -36.58 | 175.45 | 1/4/04 | full names will be added on acceptance |
| Adult | *Uropetala* | carovei | KN | Waikanae River, end of of Mangaone South Rd., 10 KM E Waikanae, 145 m | WF09 | 0.05 | 0.02 | -40.52 | 175.07 | 1/6/04 | full names will be added on acceptance |
| Nymph | *Uropetala* | carovei | KN | Waikanae River, end of of Mangaone South Rd., 10 KM E Waikanae, 145 m | WF10 | 0.05 | 0.01 | -40.52 | 175.07 | 1/27/24 | full names will be added on acceptance |
| Nymph | *Uropetala* | carovei | KN | Waikanae River, end of of Mangaone South Rd., 10 KM E Waikanae, 145 m | WF11 | 0.05 | 0.03 | -40.52 | 175.07 | 1/27/24 | full names will be added on acceptance |
| Exuvea | *Uropetala* | carovei | KS | West Coast South Island | WG01 | 0.10 | 0.06 | - | - | - | full names will be added on acceptance |
| Adult | *Uropetala* | chiltoni | KM | North tributary, Acheron River, South of Lake Lyndone, 772 M | WG02 | 0.10 | 0.06 | -43.19 | 171.4 | 1/10/04 | full names will be added on acceptance |
| Adult | *Uropetala* | chiltoni | KM | Cass | WG06 | 0.02 | 0.10 | -43.02 | 171.45 | 1/22/01 | full names will be added on acceptance |
| Adult | *Uropetala* | chiltoni | KM | Around Cass field station | WG07 | 0.07 | 0.02 | -43.02 | 171.45 | 1/22/01 | full names will be added on acceptance |
| Adult | *Uropetala* | carovei | KS | Momorangi Bay | WG08 | 0.06 | 0.07 | -41.28 | 173.94 | 12/14/00 | full names will be added on acceptance |
| Nymph | *Uropetala* | carovei | KS | Aorere Gold Fields | WG09 | 0.09 | 0.08 |  |  | 1/25/24 | full names will be added on acceptance |
| Nymph | *Uropetala* | carovei | KS | Aorere Gold Fields | WG10 | 0.10 | 0.04 |  |  | 1/26/24 | full names will be added on acceptance |
| Nymph | *Uropetala* | carovei | KS | Aorere Gold Fields | WG11 | 0.11 | 0.04 |  |  | 1/27/24 | full names will be added on acceptance |
| Nymph | *Uropetala* | chiltoni | KM | Cass | WG12 | 0.04 | 0.06 | -43.02 | 171.45 | 1/23/24 | full names will be added on acceptance |
| Nymph | *Uropetala* | chiltoni | KM | Cass | WH01 | 0.10 | 0.07 | -43.02 | 171.45 | 1/23/24 | full names will be added on acceptance |
| Nymph | *Uropetala* | chiltoni | KM | Cass | WH02 | 0.09 | 0.03 | -43.02 | 171.45 | 1/23/24 | full names will be added on acceptance |
| Exuvea | *Uropetala* | chiltoni | KM | Cass | WH03 | 0.02 | 0.20 | -43.02 | 171.45 | 1/23/24 | full names will be added on acceptance |
| Adult | *Uropetala* | chiltoni | KM | Otira valley track | WH04 | 0.03 | 0.20 | -42.9 | 171.54 | 1/19/24 | full names will be added on acceptance |
| Adult | *Uropetala* | chiltoni | KM | Otira valley track | WH07 | 0.05 | 0.25 | -42.9 | 171.54 | 1/20/24 | full names will be added on acceptance |
| Adult | *Uropetala* | chiltoni | KM | Otira valley track | WH08 | 0.02 | 0.19 | -42.9 | 171.54 | 1/21/24 | full names will be added on acceptance |
| Adult | *Uropetala* | chiltoni | KM | Otira valley track | WH09 | 0.03 | 0.17 | -42.9 | 171.54 | 1/21/24 | full names will be added on acceptance |
| Adult | *Uropetala* | chiltoni | KM | Otira valley track | WH10 | 0.03 | 0.15 | -42.9 | 171.54 | 1/21/24 | full names will be added on acceptance |
| Nymph | *Uropetala* | *chiltoni* | KM | Otira valley track | WH11 | 0.03 | 0.12 | -42.9 | 171.54 | 1/22/24 | full names will be added on acceptance |
| *Two additional KM from Cass were added from previous AHE sequencing by Tolman et al., 2024 with mean depths of 0.19 and 0.14 and missingness of 0.26 and 0.27. | | | | | | | | | | | |

| **Species** | **Raw Occurrences** | **Thinned Occurrences** | **Features** | **Regularization** | **AUCval** | **OR10** | **AICc** | **delta.AICc** | **ncoef** |
| --- | --- | --- | --- | --- | --- | --- | --- | --- | --- |
| Kapokapowai Montane | 190 | 75 | H | 3 | 0.86 | 0.09 | 983.83 | 0.43 | 8 |
| Kapokapowai North | 532 | 239 | H | 4 | 0.44 | 0.16 | 1570.48 | 1.90 | 2 |
| Kapokapowai South | 193 | 129 | LQ | 3 | 0.75 | 0.11 | 1704.57 | 1.87 | 7 |
| **Supplementary Table 2:** Sample size of occurrence localities before and after spatial thinning, and maxent ENM settings for Kapokapowai Montane, and Kapokapowai North and South for Te Ika-a-Māui and Te Waipounamu, respectively. Settings shown are feature class (features). Statistics shown are mean validation AUC (AUC_val_), mean omission rates for 10 percentile training values (OR10), Akaike Information Criterion (AICc), and delta Akaike Information Criterion (delta.AICc), and number of non-zero coefficients (ncoef). | | | | | | | | | |

| Supplementary Table 3: F_st_ values with restrictive sampling of Kapokapowai Montane | | | |
| --- | --- | --- | --- |
| Subset 1 | Putative species 2 | Mean F_ST_ | Weighted F_ST_ |
| Kapokapowai Montane (positive on PC1) | Kapokapowai South | 0.00 | 0.00 |
| Kapokapowai Montane (positive on PC1) | Kapokapowai North | 0.03 | 0.04 |
| Kapokapowai Montane (negative on PC1) | Kapokapowai South | 0.50 | 0.63 |
| Kapokapowai Montane (negative on PC1) | Kapokapowai North | 0.57 | 0.66 |

| Supplemental Table 4: Missingness of data and PC1 value of KM samples | | |
| --- | --- | --- |
| Individual | PC1 | Missingness |
| WG02 | 0.10 | 0.17 |
| WG06 | 0.07 | 0.08 |
| WG07 | 0.11 | 0.04 |
| WG12 | 0.15 | 0.02 |
| WH01 | 0.15 | 0.03 |
| WH02 | 0.10 | 0.03 |
| WH03 | -0.30 | 0.10 |
| WH04 | -0.30 | 0.13 |
| WH07 | -0.16 | 0.16 |
| WH08 | -0.31 | 0.12 |
| WH09 | -0.3 | 0.12 |
| WH10 | -0.26 | 0.07 |
| WH11 | -0.30 | 0.07 |
| Uropetala_chiltoni1* | -0.26 | 0.15 |
| Uropetala_chiltoni2 | -0.21 | 0.15 |

| Number of ancestral populations | CV error | Loglikelihood |
| --- | --- | --- |
| 1 | 1.25 | -11490.35 |
| 2 | .88 | -6232.72 |
| 3 | .90 | -5312.81 |
| 4 | .98 | -4704.61 |
| 5 | 1.1 | -4315.63 |
| Supplementary Table 5: Cross-validation error and loglikelihood assuming differing ancestral populations in the admixture analysis. | | |

| Model 1: *Argentinopetala archangelskyi* included in the ancestral state reconstruction | | | | |
| --- | --- | --- | --- | --- |
| Model | log(L) | d.f. | AIC | Weight |
| Equal-Rate | -13.18 | 1 | 28.37 | 0.66 |
| Custom ordered model | -13.18 | 2 | 30.37 | 0.24 |
| Symmetric-Rates | -13.18 | 3 | 32.37 | 0.09 |
| All-Rates-Differed | -13.18 | 6 | 38.37 | <0.01 |
| Model 2: *Argentinopetala archangelskyi* excluded from the ancestral state reconstruction | | | | |
| Equal-Rate | -12.08 | 1 | 26.17 | 0.66 |
| Custom ordered model | -12.08 | 2 | 28.17 | 0.24 |
| Symmetric-Rates | -12.08 | 3 | 30.17 | 0.09 |
| All-Rates-Differed | -12.00 | 6 | 35.98 | 0.01 |
| Supplemental Table 6: Model selection for the ancestral state reconstruction analysis with and without the fossil taxon *A. archangelskyi*. | | | | |

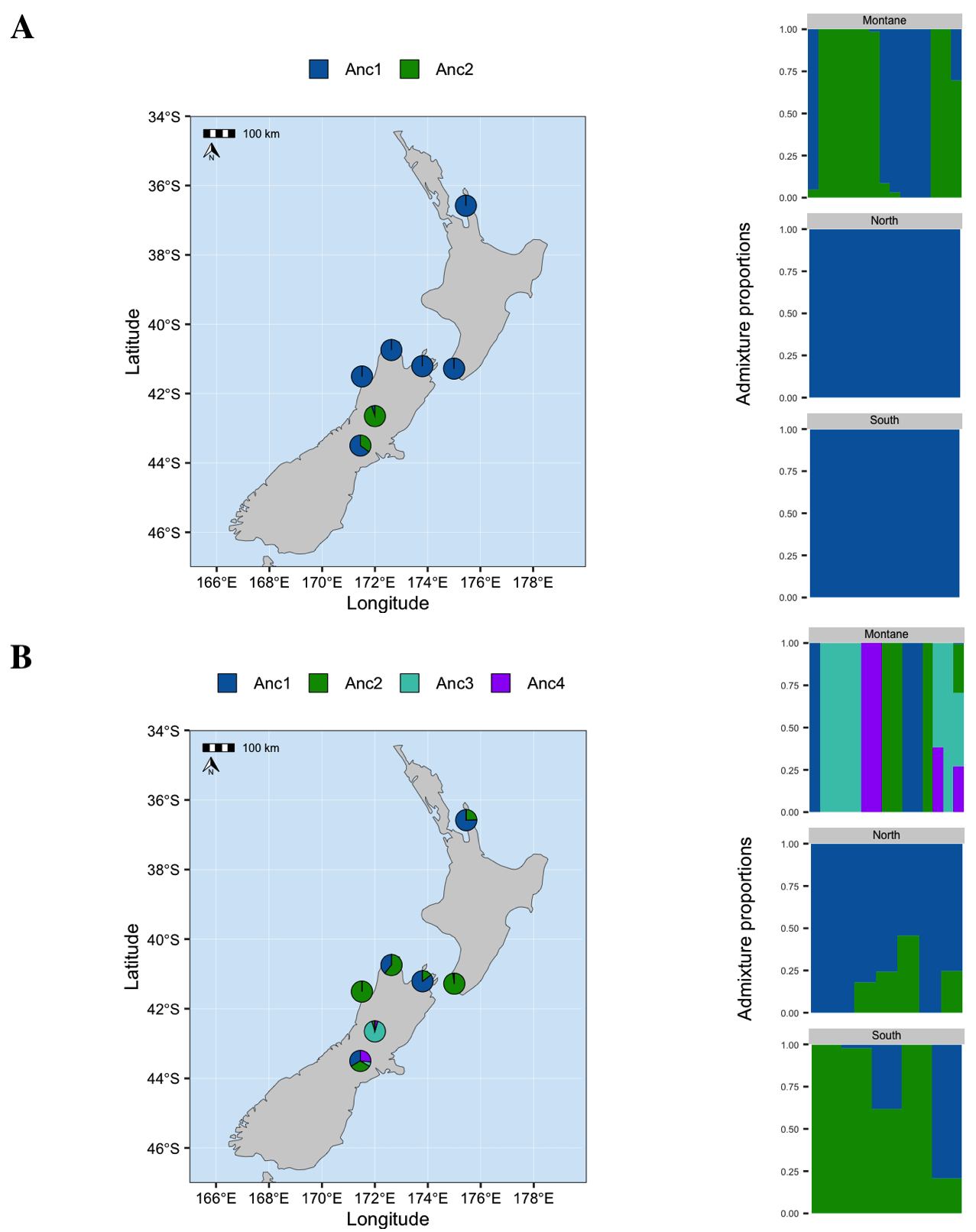

**Supplemental Figure 1:** Admixture analyses assuming (A) K=2 and (B) K=4 ancestral populations.

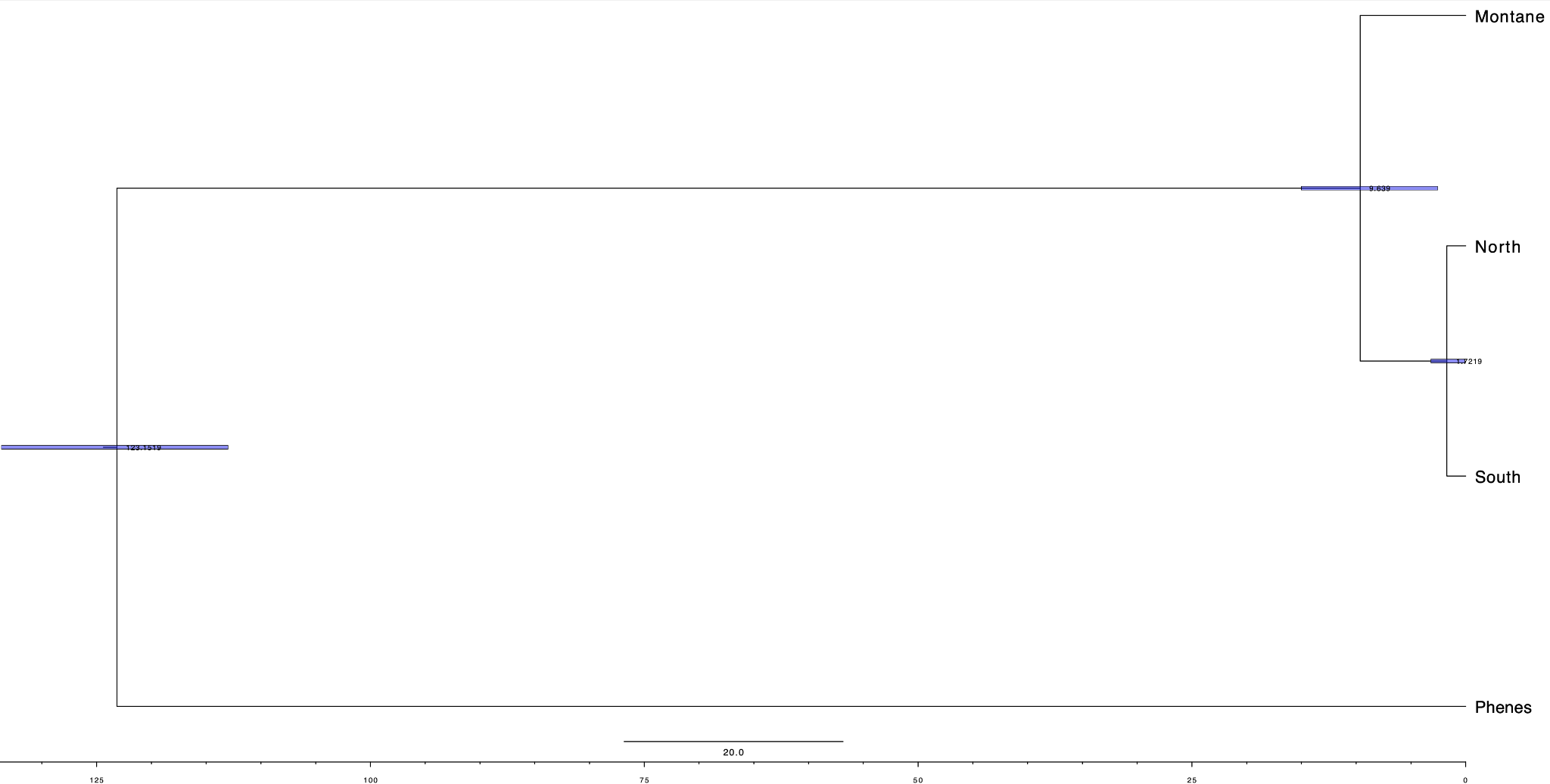

**Supplemental Figure 2:** Full dated time tree between the three populations of Kapokapowai, and the outgropu *Phenes raptor*.

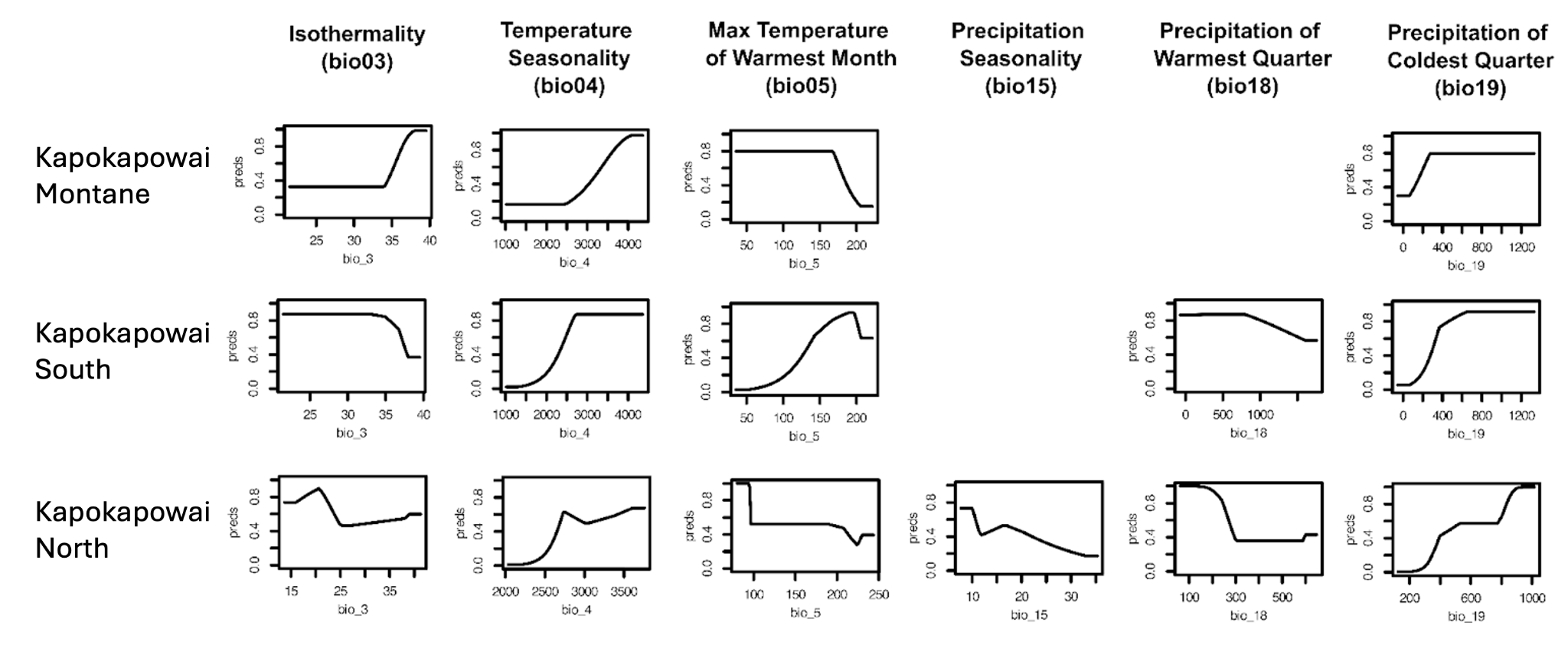

**Supplemental Figure 3:** Environmental response curves for Kapokapowai Montane and Kapokapowai South and North. Response curves show how each environmental variable individually affects the MaxEnt prediction in terms of increasing or decreasing suitability. Behavior of response curves are dictated by model complexity (feature classes and regularization multipliers).

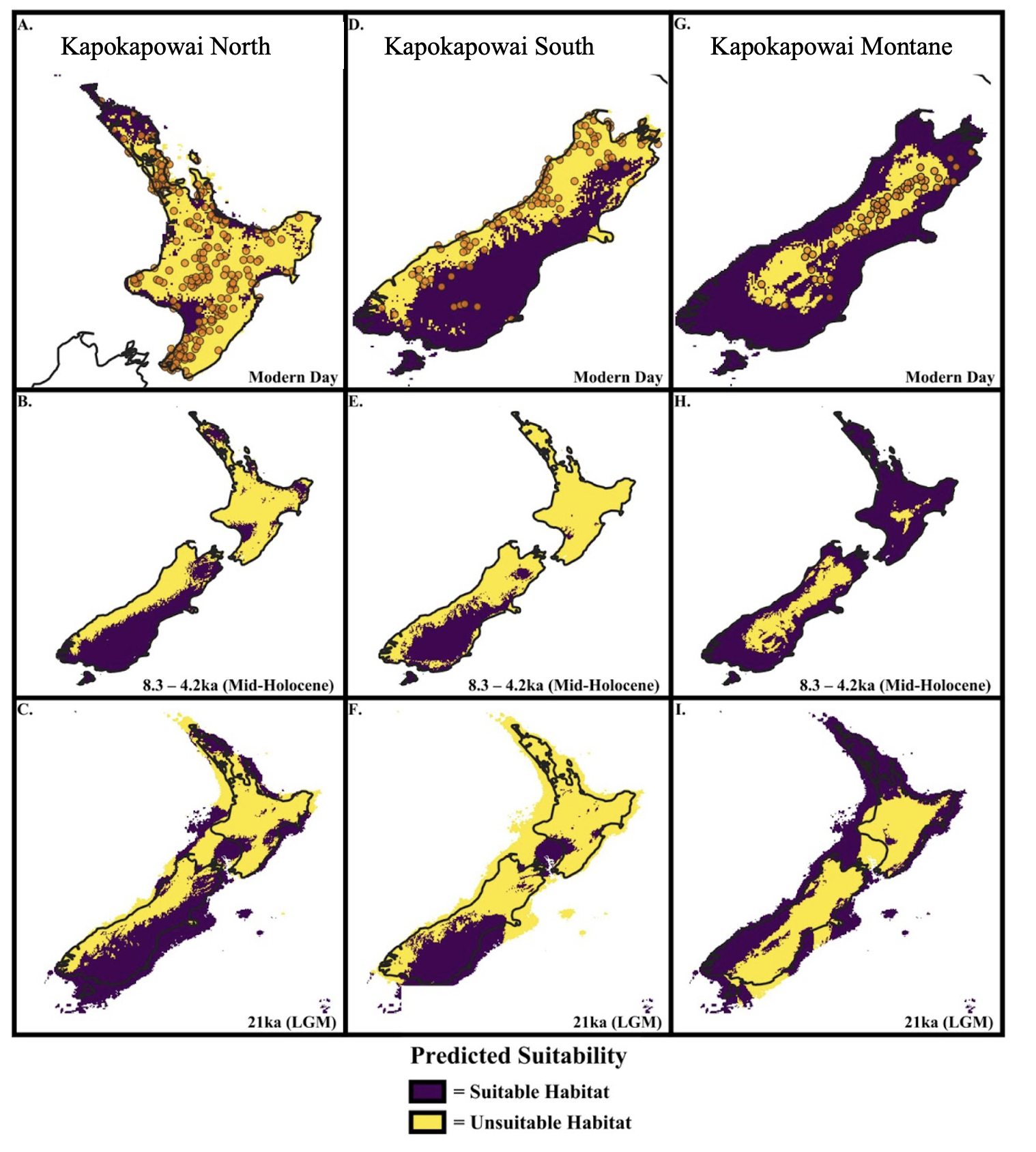

**Supplemental Figure 4:** Thresholded maxent predictions for Kapokapowai projected to the modern day using our ‘Anthropocene variables’, mid-Holocene (Northgrippian: 4.2 – 8.326ka) generated from the PaleoClim dataset (Brown et al. 2018), and Last Glacial Maximum (LGM)(21ka) and Last Interglacial (LIG)(130ka) generated from the CHELSA database (Karger et al. 2017). Predictions were derived from the optimal model using the criterion of mean valuation AUC (AUC_val_) values being the highest, and 10% omission rate being the lowest. Threshold predictions were calculated from the 10-percentile omission rate from our model evaluation. Bright colors (yellow) indicate areas deemed ‘suitable’ while colder colors (violet) indicate areas deemed ‘unsuitable’.
